# PanVA: a visual analytics tool for pangenomic variant analysis

**DOI:** 10.1101/2025.06.23.661080

**Authors:** Astrid van den Brandt, Folkert de Vries, Robin van Esch, Sander Vlugter, Dirk-Jan M. van Workum, Huub van de Wetering, Anna Vilanova, Sandra Smit

**Affiliations:** Department of Mathematics and Computer Science, Eindhoven University of Technology, Groene Loper 5, 5612 AE, Eindhoven, The Netherlands; Bioinformatics Group, Wageningen University & Research, Droevendaalsesteeg 1, 6708PB, Wageningen, The Netherlands

## Abstract

**Summary:** The growing number of sequences and increasing proof that single references create reference bias have driven the development of pangenomes to represent the genomic diversity of species. To leverage this complex diversity information for biological insights, analysis and visualization support are needed to explore the variants in the context of metadata and phylogenies. We developed PanVA, an interactive visual analytics tool for exploring sequence variants in groups of homologous sequences in their biological context. PanVA is a web application that allows users to explore existing instances or create new ones to visualize their own data.

**Availability and Implementation:** The PanVA source code is available on GitHub at https://github.com/PanBrowse/PanVA under the GPLv3 License. Documentation and and public demo instances showcasing examples can be accessed at https://panbrowse.github.io/PanVA/.

## 1. Introduction

Comparative genomics research increasingly uses pangenomes instead of single reference approaches to analyze relations between genetic variations and phenotypes. Although they allow for a more complete view of the variation, their size and complex, often graph-based, data structure hinder exploration and interpretation in the biological context (The Computational Pan-Genomics Consortium, 2018). Visualization and visual analytics are often used to explore and interpret genomics data interactively. However, existing approaches fall short in supporting pangenomic variant analysis and exploration for several reasons. First, many tools for single reference analyses cannot adequately represent the diversity of pangenomes. These include genome browsers such as IGV (Thorvaldsdóttir et al., 2013), alignment viewers such as Jalview (Waterhouse et al., 2009), and more specialized tools such as Variant View (Ferstay et al., 2013). Secondly, tools that can handle pangenomes are often tailor-made for specific (groups of) organisms, e.g., RPAN (Sun et al., 2017) and PanX (Ding et al., 2018). Lastly, the more generalized pangenome tools mainly provide views of the graph, e.g., ODGI (Guarracino et al., 2022), or higher level synteny and structural variations such as GENESPACE (Lovell et al., 2022). While these tools provide useful views of the sequences, their ability to support tasks involving sequence variant analysis and exploration is limited. These tasks require interpreting sequences alongside metadata such as phenotypes and phylogenetic information.

We developed PanVA—a visual analytics tool for interactive analysis of pangenomic variants— to overcome existing limitations. In this work, we introduce a performant, user-friendly software implementation featuring an interactive visual design (van den Brandt et al., 2023) developed and evaluated using a user-centered design approach in close collaboration with genomics experts (Sedlmair et al., 2012).

## 2. Design and Interactive Features

PanVA is an interactive visual analytics tool designed to help users perform detailed analysis of sequence variants across many gene sequences that belong to a homology group of interest (HoI). The tool interweaves views of sequence alignment data, phylogenetic information, phenotypes, and other sequence metadata, making it easier to understand the biological context of the variants. Furthermore, a variety of interactions such as selection, filtering, grouping, and aggregation are possible within this interwoven context, to further facilitate exploration and comparison tasks. For an extensive description of the supported tasks, design choices and interactions, we refer to the design study paper (van den Brandt et al., 2023). Below we briefly discuss the main views and interactive functionalities. We also detail several novelties compared to the design study prototype.

### 2.1. Views

PanVA contains two main views: the Gene Overview and the Locus view. The *Gene Overview* (Fig. 1A) shows the conservation of a selected homology group across its alignment as a compact area chart track. If gene annotations are loaded, they are shown as tracks of colored sequence segments, e.g., the coding sequences (CDS) in pink or exons in purple as shown in Fig. 1.2. The mouse pointer can be used to brush select a region of interest (RoI) for closer inspection in the Locus View. The *Locus View* ( Fig. 1B) presents four connected visualizations and interactions to display the information of the selected RoI. At the center, the selected RoI is represented as a heatmap, providing an alignment of the nucleotide sequences. Above this heatmap, a series of squares visually encode gene model annotations corresponding to the selected positions and tailored to a selected gene model. Adjacent to the heatmap, users can explore phylogenetic relationships through either phylogenetic trees or interactive hierarchical clustering dendrograms. On the opposite side, the view is organized to display phenotypes and other metadata values, which are visually encoded in various formats depending on the type of data: bar charts for quantitative metrics, circle glyphs for binary data, and colored text labels for categorical information. The collapsible side menu ( Fig. 1C) contains the Homology Table (Fig. 1D) for filtering and selecting homology groups. Furthermore, it contains legends, additional metadata about the selected homology group, and can be used to configure the views and interactions.

**Fig. 1:**
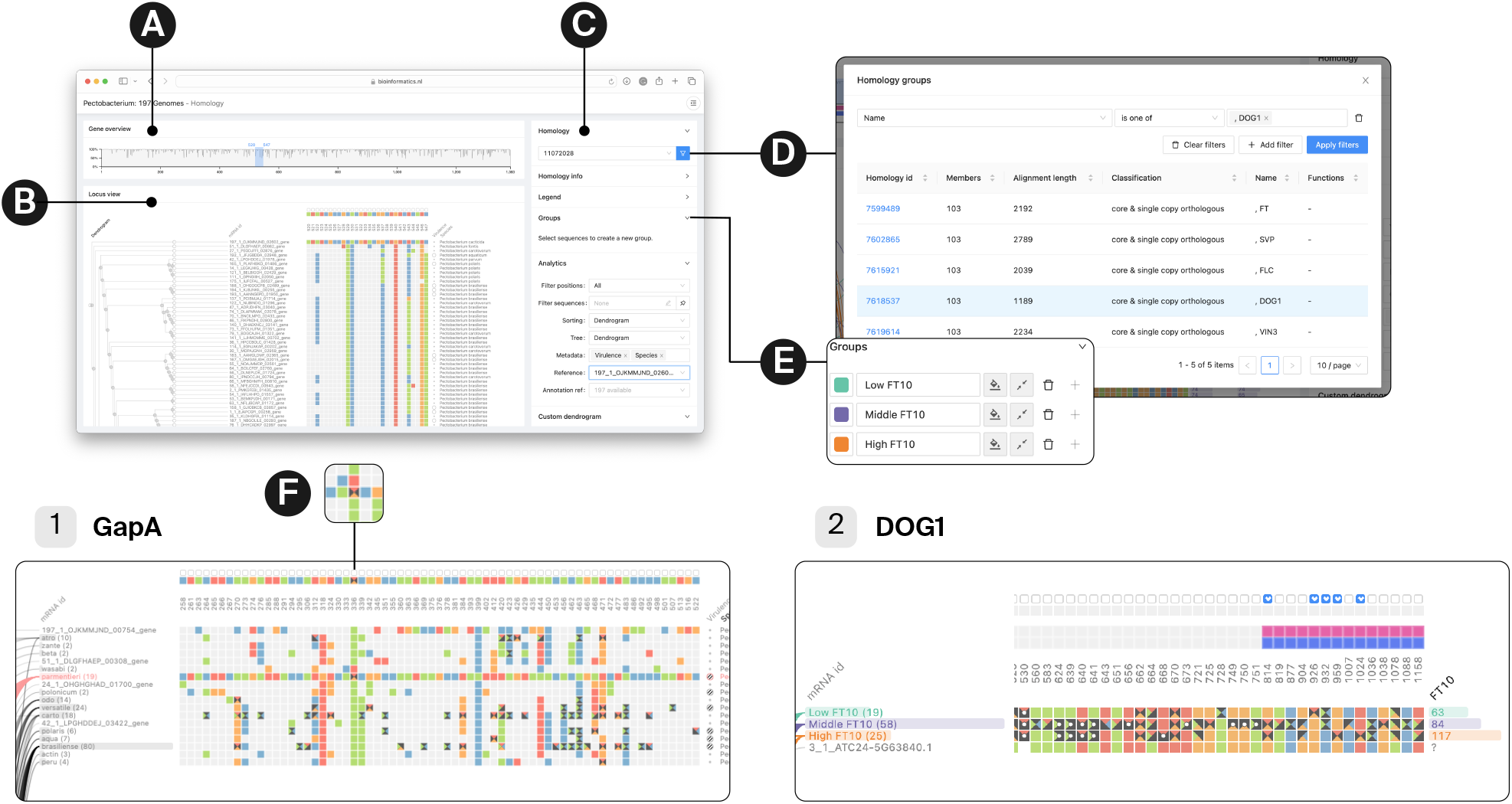
PanVA’s main interface components: (A) Gene Overview, (B) Locus View, and (C) the Control Panel. This panel supports improved selection functionality for homology groups (D) and group configuration options (E). Two use cases are demonstrated: (1) the GapA gene in *Pectobacterium*, where (F) shows the aggregation glyph, and (2) the DOG1 gene in *Arabidopsis thaliana*.

**Fig. 2:**
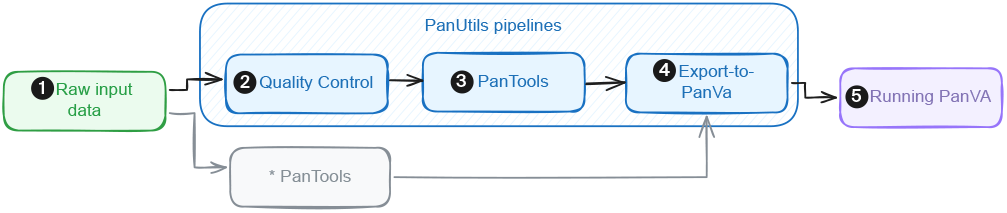
The workflow from raw input data to running PanVA instance: (1) data collection, (2) quality control, (3) necessary PanTools functions (construction, annotation, homology grouping, etc.), (4) data transformation to PanVA format, (5) PanVA instance creation. As an alternative to the snakemake pipelines (2–4), PanTools can be run standalone.

### 2.2. Interactions and Analytics

The tool supports various interactions with the data and views to support the exploration and comparison tasks effectively.

#### Data Selection and Filtering

Users can search the homology table (Fig. 1D) to find their HoI by filtering on criteria such as gene names, function (e.g., GO terms), or pangenome classification (core, accessory, unique). Filters can also be applied to the selected HoI to reduce the data shown in the visualizations of the Locus View. They can be applied on two levels: filtering entire sequences (rows) in the heatmap and metadata visualizations or alignment positions in the heatmap only (columns). Available position and sequence metadata features can be used for filtering. Users can save the filter settings to apply them to the next selected HoI. Lastly, any selected sequence or group is sequences is highlighted in corresponding visualizations. For groups, the highlight color can be customized and the user can toggle the visibility.

#### Sorting and Clustering

The information of the Locus View can be sorted in various ways. By default, the selected tree or hierarchical clustering dendrogram determines the row order of the heatmap and metadata. Users can change the ordering in thee ways: by changing the tree type, by a selected metadata feature, or by one ore more positions in the alignment. In case multiple positions are selected (which can be a range or discontinuous set of positions), a custom clustering result is computed and the corresponding dendrogram is shown. Users can choose to display this new dendrogram or any other tree, which is referred to as *linking* or *unlinking* in the tool. When rows are sorted or trees are *unlinked*, lines between the dendrogram and heatmap connect the corresponding rows. During sorting and clustering, transitions are animated to avoid confusion in the exploration process and users can maintain a mental map. However, with a large RoI selected the transitions might become slow. Therefore, transitions can be switched on and off from the side menu.

#### Grouping and Aggregation

Sequences and corresponding metadata can be grouped through mouse selection and can be further configured from the side menu (Fig. 1E). By default, sequences in groups are collapsed into one aggregated row showing a custom aggregation glyph for positions containing multiple nucleotide types, for example in Fig. 1F sequences in the pink highlighted group either have an A (red) or T (orange) on position 336. Groups can be shown collapsed or expanded, with a custom highlight color and name.

### 2.3. Novel Features

There are four novel interactive features and views of the tool compared to the design study prototype. In terms of selection and filtering interactions, the tool now presents a homology search and query functionality, options to filter sequences (rows) in the Locus View, and the ability to save filter settings for other homology groups. Furthermore, the annotation view is extended with options to visualize multiple gene model annotation features, such as exon and intron segments in addition to coding sequences. Beyond these extensions, the implementation has been transformed from proof-of-concept design prototype to a fully functional tool that can be used in genomics research.

## 3. Implementation

PanVA is implemented as a web-based application, using modern Javascript libraries to handle visualization and interactions for exploring large-scale pangenomic data. The Canvas API is used to optimize the rendering of large RoIs from the multiple sequence alignment with optional animated transitions. Users can customize the tool according to their needs using a runtime configurable JSON specification that provides dynamic metadata support. The online documentation provides an example JSON specification file for initial setup. Users can further customize and modify graphics and interactions directly from the tool’s side menu, e.g., high DPI rendering, custom group names and color palettes.

PanVA can be accessed in various ways. Demo instances are publicly available to try out the tool’s functionalities for a variety of pangenomes. Users can run PanVA locally on their own data with docker or directly from the source code to create their own instances. Resources and instructions on how to run PanVA are available at our GitHub repository (https://github.com/PanBrowse/PanVA).

PanVA takes preprocessed alignments, metadata tables, and tree calculations per homology group as input. The data input format is specified in our GitHub repository. We use PanTools (Jonkheer et al., 2022) to retrieve these homology groups and their corresponding metadata and phylogenetic features. The entire workflow from input data to PanVA instance, entails several steps, see Fig. 2: (1) downloading and organizing raw data, (2) performing quality control, (3) running the necessary PanTools functions, (4) converting homology groups and related metadata to PanVA input, and (5) launching the PanVA docker instance. Detailed instructions on the entire process from raw data to PanVA instance can be found in the online documentation of PanVA (https://panbrowse.github.io/PanVA/) and PanTools (https://pantools.readthedocs.io).

## 4. Use Cases

We demonstrate PanVA with bacterial and plant pangenome use cases. The demo instances linked to these use cases can be accessed through our online documentation.

### 4.1. Pectobacterium

We constructed a pangenome of 197 annotated *Pectobacterium* genomes with, amongst others, Virulence and Species as metadata to investigate genetic variation underlying virulence and host specificity (Jonkheer et al., 2021) and preprocessed the data for PanVA. Using the homology table in PanVA, we searched and filtered for genes of interest. We selected the housekeeping gene gapA, a commonly used marker gene (Cigna et al., 2017), and explore species-specific variations for *P. parmentieri*, a small and uniform species. We sorted and grouped the data on Species, filtered for variable positions, and set the *P. parmentieri* group as a visual reference for comparison. To create a compact view, we aggregated all species groups except the *P. parmentieri* group. Browsing through this view (Fig. 1.1), we found a SNP at position 399 to distinguish *P. parmentieri* from the other species.

Applying the same exploration strategy for the more variable housekeeping gene GyrB and its variations for *P. brasiliense* enabled us to discover six positions where all virulent accessions share a particular SNP pattern: 615 A, 885 T, 939 C, 1032 T, 2350 T and 2517 C, potentially suggesting this genotype is required to be virulent. However, also two avirulent strains and ten strains with an unknown phenotype share this exact pattern. Therefore, next steps would be to investigate this pattern in other housekeeping genes and collect more phenotypes for the unknown strains to further explore this discovery.

### 4.2. Arabidopsis thaliana

To analyze genetic variations relating to flowering time in *Arabidopsis thaliana*, we built a pangenome of 25 assembled genomes with 19 resequenced accessions and added flowering time phenotypes FT10 and FT16 (days to flowering time at 10 and 16 degrees Celcius). One of the genes with a known association with flowering time is DOG1 (Alonso-Blanco et al., 2016). We selected the homology group containing this gene via the Homology Table. Once loaded, we filtered, sorted, and created four groups (low FT10, high FT10, middle values, and unknown) to explore potential genetic variations for FT values (Fig. 1.2). From the aggregated group sequences, we spotted that group “ low FT” is more conserved than the others. We check mark 14 positions inside CDS regions that could be candidate SNPs and use those positions for the custom dendrogram to further check the similarity. Interactively exploring—i.e., expanding and collapsing groups and highlighting bipartite links—revealed two clusters of similar accessions (on the selected 14 positions) with very high flowering times.

## 5. Conclusion

We presented PanVA, a web-based visual analysis tool designed for exploring pangenomic variants. This tool offers a dynamic interface that allows users to analyze sequence variants within the context of metadata and phylogenetic data. PanVA visually integrates genomic data and metadata, and supports exploration of diverse organisms and pangenomes. We demonstated the versatility of our tool with use cases of 197 *Pectobacterium* genomes and 3 *Arabidopsis thaliana* genomes with 100 resequenced accessions.

Compared to the design study, we addressed some shortcomings, such as displaying the degenerate bases and filtering genomes. However, other limitations remain, such as the reliance on preprocessed data rather than a live connection to the pangenome graph. Furthermore, flexible data selection (i.e., including flanking sequences) would be helpful to analyze regulatory regions. Looking ahead, we plan to enhance PanVA by adding views for exploration of larger regions encompassing multiple homology groups or gene sets in separate areas. Finally, visual scalability could be improved to increase information density. These extensions will further enhance the tool’s capability for exploratory analysis variants in pangenomes.

